# Synthesis and evaluation of novel copper-antibody conjugates for the chemodynamic therapy of HER2-positive breast cancer

**DOI:** 10.64898/2026.04.30.721915

**Authors:** Daria Otvodnikova, Kirill Chernov, Sofia Gornostaeva, Maksim Meshechko, Oleg Kuchur, Vladimir Sharoyko, Sergey Tsymbal

## Abstract

In this work we present antibody-metal conjugate as a new subclass of antibody-drug conjugates (ADC) for the chemodynamic therapy of cancer based on the rapid generation of reactive oxygen species (ROS) upon copper reduction. We used conventional therapeutic antibody trastuzumab and DOTA-NHS ester for the design and initial proof-of-concept. Thus, trastuzumab-DOTA-copper conjugate (TDCC) was synthesized. We demonstrate that TDCC retains specific binding to HER2-positive cancer cells with approximately native immunoreactivity and achieves stable copper incorporation with an average drug-to-antibody ratio of up to ∼8. In the presence of physiological reducing agents such as N-acetylcysteine or cysteine, TDCC generates substantial reactive oxygen species (ROS), leading to pronounced cytotoxicity and long-term suppression of clonogenic survival in HER2-positive SK-BR-3 and BT-474 cells. Notably, HER2-negative MDA-MB-231 cells and non-malignant HS5 fibroblasts remain largely unaffected, confirming target-dependent activity. The conjugate remains stable under storage conditions for up to 30 days, and the DOTA linker itself does not interfere with copper-mediated redox chemistry. Our findings identify TDCC as a novel class of targeted oxidative stress inducers that exploit the vulnerability of HER2–positive tumors to copper–mediated cytotoxicity. This strategy not only preserves the specificity of antibody–based delivery but also introduces a distinct mechanism of action capable of bypassing conventional resistance pathways, warranting further preclinical development.

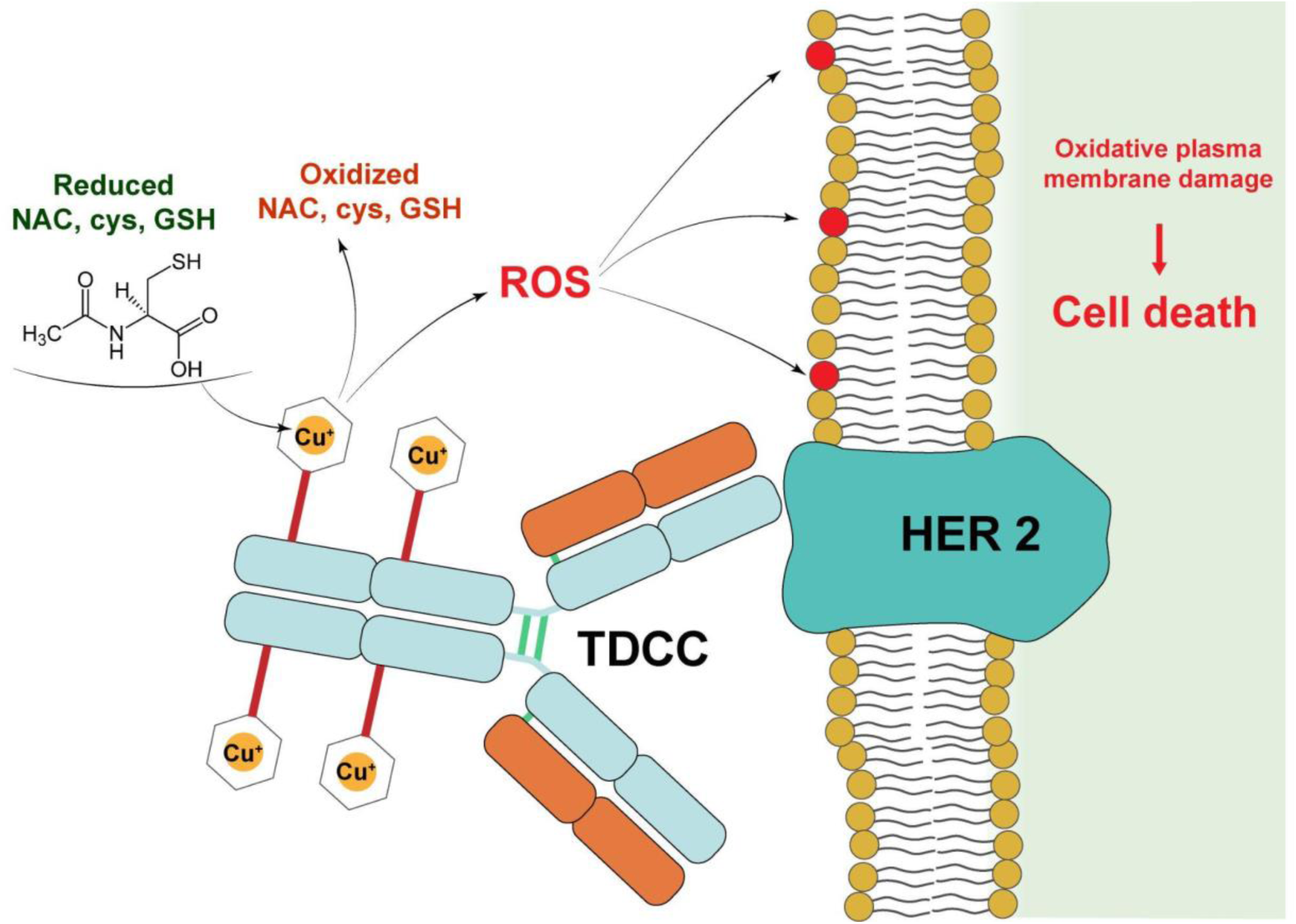

## 1. Introduction

The landscape of modern anticancer therapy has been transformed by remarkable advances over the past two decades: 5-year survival rates for many malignancies have soared, and diseases once considered incurable are now manageable with long-term treatment strategies [1]. A significant breakthrough was the advent of immunotherapy, a method using monoclonal antibodies that target specific molecular markers expressed on cancer cells. Trastuzumab (Herceptin), cetuximab, nivolumab, and many other immunotherapeutic drugs are now reliable agents in a variety of solid tumors and hematologic malignancies, demonstrating both efficacy and acceptable safety profiles [2,3]. Particularly, trastuzumab has substantially improved outcomes for HER2-positive breast cancer patients, yet acquired resistance remains a major clinical hurdle, necessitating strategies with distinct mechanisms of action [4,5]. This clinical success has profoundly influenced the pharmaceutical industry, redirecting substantial investment from traditional small-molecule inhibitors to the development of biopharmaceuticals, including antibody-drug conjugates (ADCs) and immune checkpoint inhibitors [6,7]. ADCs represent a logical evolution of monoclonal antibody therapy: by linking highly potent payloads to tumor-targeting antibodies, they combine the specificity of immunotherapy with the potency of conventional chemotherapy [8]. Several ADCs, such as trastuzumab emtansine (T-DM1) and trastuzumab deruxtecan (T-DXd), have already transformed the treatment landscape for HER2-positive breast cancer, particularly in patients who progress on trastuzumab-based regimens [9,10]. With targeted therapy achieving unprecedented specificity, the focus has shifted toward overcoming the biological complexity of tumors. While the initial response to targeted agents like trastuzumab can be dramatic, a significant proportion of patients eventually develop resistance [11]. In HER2-positive breast cancer, which accounts for approximately 15-20% of all cases, resistance to trastuzumab arises through multiple mechanisms, including the upregulation of alternative signaling pathways (PI3K/AKT/mTOR), epitope masking, and the selection of cancer stem-like cells [4,12,13]. This adaptive capacity of tumors represents the main challenge for modern cancer therapy: drug resistance, both intrinsic and acquired, ultimately limits the durability of clinical responses and necessitates the development of mechanistically distinct approaches [14,15].

Copper ions, while capable of triggering oxidative cell death via Fenton–like chemistry, lack tumor selectivity, precluding their systemic use. In our previous work we demonstrated that copper complexes can induce potent cytotoxicity in cancer cells through redox cycling in the presence of physiological reducing agents such as N–acetylcysteine (NAC) or glutathione [16,17]. However, to harness this cytotoxic potential while ensuring tumor specificity, we hypothesized that conjugating copper ions to a tumor–targeting antibody could provide a solution. Trastuzumab, with its well–characterized binding to the extracellular domain of HER2, serves as an ideal delivery vehicle. By attaching copper via a stable chelator, we aimed to create a construct that remains inert during circulation but becomes actively cytotoxic upon reduction by exogenously added thiols such as NAC or cysteine, generating a localized oxidative burst capable of eradicating HER2–positive tumor cells, including those with acquired resistance. For this purpose, we selected DOTA (1,4,7,10–tetraazacyclododecane–1,4,7,10–tetraacetic acid), a macrocyclic chelator widely used in radiopharmaceuticals for its high affinity and kinetic stability with various metals, including copper [18,19]. The DOTA moiety can be readily coupled to lysine residues of the antibody via an N–hydroxysuccinimide (NHS) ester, enabling reproducible conjugation without compromising the antibody’s binding integrity [20,21].

In the present study, we report the synthesis, characterization, and in vitro evaluation of a trastuzumab–DOTA–copper conjugate (TDCC). We systematically assessed its stability under different storage conditions, its ability to retain specific binding to HER2-positive cancer cells, and its capacity to generate reactive oxygen species (ROS) in the presence of the physiological reductants such as NAC or cysteine. Furthermore, we compared its cytotoxicity on HER2-positive (SK-BR-3, BT-474) and HER2-negative (MDA-MB-231) breast cancer cell lines, as well as non-malignant human fibroblasts HS5, to evaluate selectivity and therapeutic potential. Our findings demonstrate that TDCC combines the targeting precision of trastuzumab with the pro-oxidant cytotoxicity of copper, offering a novel strategy for the treatment of HER2-positive breast cancer, particularly in settings where resistance to conventional therapies has emerged.

## 2. Materials and Methods

### 2.1 Antibodies and chemicals

Sodium hydrocarbonate (130272), LenReactiv, Saint Petersburg, Russia

Sodium carbonate (130274), LenReactiv, Saint Petersburg, Russia

Milli-Q grade water prepared with ICW-3000, Merck Millipore, Burlington, MA, USA

DOTA-NHS ester (4245-500mg), Lumiprobe Rus, Moscow, Russia

Trastuzumab liofilizate, Ozon Pharmaceuticals, Togliatti, Russia

Bradford Reagent (ML106), Himedia Laboratories, Thane, India

PBS tablets (18912014), Thermo Fisher Scientific, Uppsala, Sweden

Acrylamide 2x crystallized (Х122), PanEco, Moscow, Russia

N,N′-Methylenebisacrylamide (Х160), PanEco, Moscow, Russia

Tris-OH (H-T6791-0.5), Helicon, Moscow, Russia

TRIS hydrochloride (RES3098T-B701X), Sigma-Aldrich, Saint Louis, MO, USA

TRIS 1.0 M buffer pH 6.8 (J63831), Alfa Aesar, Ward Hill, MA, USA

Tris-OH BioXtra (H-T6791-0.5), Sigma-Aldrich, Saint Louis, MO, USA

Glycine pure EP (LC-10273.1000), NeoFroxx, Einhausen, Germany

2-Mercaptoethanol (M6250), Sigma-Aldrich, Saint Louis, MO, USA

Glycerol for molecular biology ≥99% (G5516), Sigma-Aldrich, Saint Louis, MO, USA

Bromophenol Blue (114391), Sigma-Aldrich, Saint Louis, MO, USA

Sodium dodecyl sulfate (CR2209004), Servicebio, Wuhan, PRC

Tween® 20 for molecular biology (A4974), AppliChem, Darmstadt, Germany

Acetic acid (64-19-7), Ekos-1, Moscow, Russia

Coomassie G-250 (О120), PanEco, Moscow, Russia

TEMED (17131201), GE Healthcare, Chicago, IL, USA

Ammonium persulfate (A3678), Sigma-Aldrich, Saint Louis, MO, USA

Rav-11 protein marker (PS-2050), Biolabmix, Novosibirsk, Russia

Copper(II) chloride dihydrate (120215), LenReactiv, Saint Petersburg, Russia

L-arginine (74-79-3), Reanal, Budapest, Hungary

Sodium acetate 3-hydrate (130883, LenReactiv), Macco Organiques, Bruntál, Czech Republic

Sodium hydroxide (130108), LenReactiv, Saint Petersburg, Russia

Sodium Chloride (130314), LenReactiv, Saint Petersburg, Russia

Potassium Chloride (100244), LenReactiv, Saint Petersburg, Russia

Disodium Phosphate (120298), LenReactiv, Saint Petersburg, Russia

Monopotassium Phosphate (100230), LenReactiv, Saint Petersburg, Russia

Hydrochloric acid (170266), LenReactiv, Saint Petersburg, Russia

Fluorescein 5(6)-isothiocyanate (46950), Sigma-Aldrich, Saint Louis, MO, USA

Dimethyl sulfoxide 99.9%, Ekos-1, Moscow, Russia

Conjugate of mouse mAb against human Fc with HRP, Hema, Saint Petersburg, Russia

Recombinant HER2 Fc-tag (HE2-H5253-100ug), ACROBiosystems, Newark, DE, USA

Protein A (R060), Hema, Saint Petersburg, Russia

Casein fat free (C 1686), Ottokemi, Mumbai, India

D(+)-Sucrose 99.7% (17714-, ChemMed Group), Acros organics, Geel, Belgium

Tetramethylbenzidine (R055), Hema, Saint Petersburg, Russia

### 2.2 Synthesis of Trastuzumab-DOTA-Copper conjugates (TDCC)

A purified solution of trastuzumab (M = 145.53 Da), dissolved in sodium bicarbonate buffer (0.2 M, pH = 9.2), to a final antibody concentration of 4-5 mg/mL was mixed with DOTA-NHS ester (M_r_ = 761.49 g/mol), prepared at a 120-fold molar excess in the same buffer, followed by gentle mixing through pipetting twenty times. The mixture was carefully shaken and incubated overnight at +4°C on a magnetic stirrer [22–24]. Unbound DOTA-NHS ester was removed using either 1) size-exclusion chromatography (SEC) or 2) dialysis. To the purified DOTA-trastuzumab conjugate, copper(II) chloride (CuCl₂) salt was added at a 10-fold molar excess over the antibody in sodium acetate buffer (0.1 M, pH 5.6) and incubated for 30 minutes on a magnetic stirrer at room temperature [25,26]. The product was then purified and, if necessary, concentrated by lyophilization (0.25 mbar, -60°C, 2-4 hours).

### 2.3 Purification methods

As was mentioned above, purification of antibodies was performed with either SEC or dialysis, depending on the processing scheme. PD-10 column loaded with Sephadex-G25 (17085101, Cytiva, Marlborough, MA, USA) used for small volume purification (less than 5 mg of a protein) of the processing fractions. The column was washed three times with Milli-Q water, followed by three washes with either sodium bicarbonate buffer or PBS, depending on the synthesis step. Liquid flow was run by gravitational force. Fractions of 1 mL were manually collected into 1.5 mL microtubes (Jet Biofil, Guangzhou, PRC). These fractions were characterized with the optical density measurement on 280 and 290 nm wavelengths using Implen N50 droplet spectrophotometer (Implen, Darmstadt, Germany). The purified protein was obtained out of 3-5 fractions poured together depending on the maximum intensity of optical density measurements.

Dialysis of the sample volume less than 10 mL was used as an alternative method of purification and a desalting step to dispose of unbound DOTA-NHS and copper salt in the reaction mixture. A dialysis membrane with 14 kDa molecular cutoff (MC30 Membra-Cel, Viskase, Lombard, IL, USA) was rinsed 15 min by Milli-Q water, then boiled for 20 min in water solution of 10 mM Na_2_EDTA and 10 mM Na_2_CO_3_. 10 cm of the membrane was clamped and checked for leakage. The sample was placed in the dialysis sack and clamped on the other end with 1-2 cm of air above the liquid level. A one-liter beaker, poured with 100-times of a sample volume of sodium acetate buffer pH = 5.6, the closed dialysis tubing was placed in the beaker and left with a magnetic stirrer for 24-36 h mixing at +4°C. The buffer was changed 3-4 times. Upon the dialysis end, the upper tubing clamp was removed and the liquid sample was taken with an autopipette. The dialysis membrane then was stored in 20% ethanol at +4°C for further uses.

### 2.4 Gel-electrophoresis

The stability, degradation and presence of light and heavy chains for antibody and conjugates were assessed by denaturing gel electrophoresis under reducing and non-reducing conditions. The 8% resolving and 5% stacking gels were prepared with 30% AA:BA acrylamide mixture, 1 M Tris-OH buffer pH 6.8 – for stacking gel, 1,5 M Tris-OH buffer pH 8.8 – for resolving gel, 10% SDS solution and deionised water, followed by addition of 100 µL 10% APS and 10 µL TEMED, and immediate casting. The running buffer was 1x TGB with 0.1% SDS. Protein samples of 20-25 μg were mixed with 6x Laemmli loading buffer, with and without β-mercaptoethanol, heated at 95°C for 5 min, injected, and run at 120 V for 100 min. Rav-11 protein marker was used for molecular weight assessment.

Upon a run ending, the gel was fixed at 40% ethanol and 10% acetate for 30 min, stained overnight with 0.1% Coomassie G-250, and destained at 20% ethanol and 10% acetate until achieving transparency. Gel imaging was done on the ChemiDoc Touch (Bio-Rad, Hercules, CA, USA) imaging system on a transparent tray, using Stain Free Gel registration with a 7.5 sec exposure. Band size and intensity were annotated manually using Image Lab Bio-Rad version 6.0 software (Bio-Rad, Hercules, CA, USA).

### 2.5 Indirect non-competitive enzyme-linked immunosorbent assay (ELISA)

Protein A was dissolved in 5 µg/mL in binding buffer (pH 9.6) containing 0.1 M sodium carbonate and added in volume 100 µL per well and incubated at +37 °C with shaking (600 rpm) for 1 h in thermal shaker PST-60HL-4 (BioSan, Latvia). After removal the content of wells, plate was blocked with 200 µL of blocking buffer containing 1% of casein, 0.015 M of NaCl, 5% of sucrose (ChimMed Group, Russia), 0.005 M potassium di-hydrogen phosphate (PanReac Quimica, Spain), 0.05% ProClin 300 (Sigma Aldrich, Germany) and incubated at +37 °C with shaking (600 rpm) for 1 h. Recombinant HER2 Fc-tag (AcroBiosystmes, USA) antigen was diluted in dilution buffer 1% of casein, 0.005 M potassium di-hydrogen phosphate, 0.05% ProClin 300, 0.25 M of tris(hydroxymethyl)aminomethane, 0.15 M sodium chloride, 0.25% Tween-20, 0.24 M hydrochloric acid, 10 % of porcine blood serum to a concentration of 5 µg/mL, and then it was serially diluted 3-fold to final concentration and added in a volume of 100 µL per well, followed by 1 h incubation at 37 °C. Plates were washed three times by adding 300 µl with a washing buffer containing 0.25 M tris(hydroxymethyl)aminomethane, 0.15 M sodium chloride, 0.25% Tween-20, and 0.24 M hydrochloric acid.

Samples were diluted to 5 µg/mL in a dilution buffer and added (100 µL per well) and incubated for 1 h at +37 °C, followed by three washing steps. Anti-human Fc monoclonal antibody conjugated with horseradish peroxidase, were sequentially diluted 1:10000 and added in volume of 100 µL per well, incubated for 1 h at 37 °C, and washed five times. Tetramethylbenzidine was added in 100 µL volume and incubated for 15 min in the dark at room temperature. The reaction was stopped by adding 100 µL of stop solution, and optical density was measured at 450 nm using a microplate reader Epoch (BioTek Instruments, Winooski, VT, USA).

### 2.6 Atomic emission spectroscopy

Arc-induced atomic emission spectroscopy (AES) was used to evaluate the amount of copper in the conjugate [27]. The sample and calibration solutions of copper(II) chloride were added into the craters of graphite electrodes, pre-filled with compacted graphite powder, in a volume of 35 µL each. The electrodes with the samples were then dried in an oven for 15 minutes at 60 °C. Subsequently, evaporation and atomization of the samples occurred in the electric arc plasma of an Iskroline spectrometer (Iskroline, Russia). A two-step operating mode was used: Step 1 - current of 18 A for 2 s, Step 2 - current of 18 A for 60 s, with the following registration parameters: step time 35 ms, frame time 0.12922 s, number of frames 400, decimation factor 1, and a total exposure time of 51.688 s. The copper concentration was determined by the emission intensity at the spectral lines of 324 nm and 327 nm using the calibration curve. The 324 nm characteristic line proved to be the most relevant.

For concentration calculation linear calibration curve equation was used, with R^2^ coefficient more than 0,9 determining the relevance of thе curve.

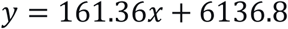

where

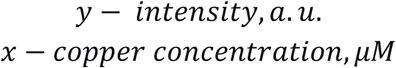

In order to calculate DAR, the following formula was used:

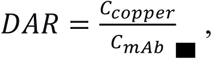

where

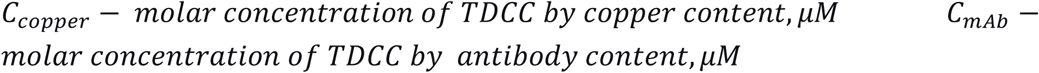

### 2.7 Protein Quantification

Protein concentration was determined using the Bradford assay (Bio-Rad Protein Assay – Standard Procedure for Microtiter Plates, Bio-Rad, USA). Optical absorbance was measured at 570 nm; a calibration curve was generated using the BSA (Biolot, Russia) or trastuzumab as standards. The protein concentration in experimental samples, along with the standard deviation, was determined by interpolating their ratios from this curve.

### 2.8 Characterization of copper-DOTA binding

Binding of copper by DOTA-chelator was confirmed spectrophotometrically by measuring optical density of copper phosphate that forms in a 1x PBS solution [28,29]. The experiment procedure started with copper chloride solution (150 mM, pH=7, in Water) mixed with 1x PBS (Sigma Aldrich, Germany) in various ratios (from 5:1 to 1:5) with total volume in a 96-well plate 100 µL per well. Simultaneously the same dilutions were prepared using the same reagents, but with DOTA dissolved in 1x PBS (150 mM). All spectra were recorded at 450 nm using Tecan Spark (Tecan, Switzerland) equipment. Obtained data were analyzed and processed graphs built in Microsoft Excel software.

### 2.9 Synthesis of Trastuzumab-FITC conjugate

All procedures were performed under cold conditions and protected from light. Prior to conjugation, trastuzumab (2 mg/mL) was transferred into 0.2 M sodium carbonate-bicarbonate buffer (pH 9.0) using a PD-10 Desalting Column packed with Sephadex G-25 (Cytiva, USA) to remove low-molecular-weight components and to provide alkaline, amine-free conditions optimal for FITC coupling to lysine residues. Eluted fractions (1 mL) were collected, and protein-containing fractions were identified by measuring absorbance at 280 nm using an Implen NanoPhotometer N60/N50 (Implen, Germany). Fractions with the highest protein concentration were pooled for subsequent labeling. FITC was freshly dissolved in DMSO to a stock concentration of 50 mM and added to the antibody solution at a molar antibody:FITC ratio of 1:16. The reaction mixture was incubated overnight at 4 °C in the dark. Unbound FITC was removed by gel filtration on a PD-10 Desalting Column (Cytiva, USA) equilibrated with PBS (pH 7.4). Eluted fractions (1 mL) were collected and analyzed spectrophotometrically at 280 and 495 nm to determine the degree of labeling (DOL) using the following equation:

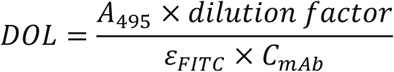

where

𝐴_495_ is the absorbance of the conjugate at 495 nm,

𝜀_𝐹𝐼𝑇𝐶_ is the molar extinction coefficient of FITC (0.69 × 10^5^𝑀^−1^𝑐𝑚^−1^),

𝐶_𝑚𝐴𝑏_ is the molar concentration of the antibody.

The antibody concentration was corrected for FITC absorbance at 280 nm using the following equation:

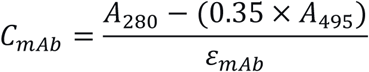

where

0.35 is the correction factor for FITC absorbance at 280 nm,

𝜀_𝑚𝐴𝑏_ is the molar extinction coefficient of trastuzumab (2.1 × 10^5^ 𝑀^−1^𝑐𝑚^−1^)

The purified FITC-conjugate was stored in PBS at 4 °C in the dark for up to 30 days.

### 2.10 Cell Lines Culturing

SK-BR-3 and BT-474 HER2-positive breast carcinoma cell lines were cultured in RPMI-1640 media (Biolot, Russia) with 10 μg/mL of insulin added specifically for BT-474. The HS-5 fibroblast cell line, cultured in the same medium, served as a non-malignant control. The MDA-MB-231 triple-negative breast cancer cell line was utilized as HER2 non-expressing model, cultured in Dulbecco’s modified Eagle’s medium (DMEM, Biolot, Russia). Each cell line was supplemented with 10% fetal bovine serum (Biolot, 1.1.8.9.), penicillin G and streptomycin (Biolot, 1.3.18.) at concentrations of 100 U/mL and 100 μg/mL, respectively, and incubated at 37 °C, 5% CO2 in a humidified atmosphere. Adherent cells were detached with trypsin-Versen (Biolot, 1.2.2.5, 1.2.3.2) mixture or Versen solution alone. All cell lines were purchased from the American Type Culture Collection (Manassas, VA, USA).

### 2.11 Flow Cytofluorometry

#### 2.11.1 ROS formation

Assessment of reactive oxygen species (ROS) accumulation by TDCC in combination with NAC was performed using the fluorescent dye 2′,7′-dichlorodihydrofluorescein diacetate (H₂DCFDA) by flow cytofluorometry. SK-BR-3 and MDA-MB-231 cells were seeded at a density of 1 × 10⁵ cells and incubated overnight to adhere. The next day, TDCC and copper chloride were added at a concentration of 5 µM, and NAC at 1 mM, and the cells were incubated for 4 hours. Positive controls included hydrogen peroxide at 1 mM added 1 hour before the end of incubation, as well as the combination of TDCC and NAC incubated for 24 hours. Cells were washed once with warm Hanks’ balanced salt solution and then incubated for 40 minutes at 37 °C with 10 µM H₂DCFDA prepared in serum–free RPMI–1640 medium or Hanks’ solution. After the dye incubation, cells were washed again with complete RPMI-1640 medium, held for 10 minutes, detached, and centrifuged for 5 minutes at 300g. The resulting pellet was resuspended in 200 µL of PBS (pH 7.4). Fluorescence of the cell suspension was analyzed on a Cytoflex flow cytometer using a blue laser (488 nm) and the FITC channel (525/40 nm).

#### 2.11.2 Competitive binding

SK-BR-3 (HER2-positive) and MDA-MB-231 (HER2-negative) cells were detached from culture dishes and transferred to microcentrifuge tubes at a density of 1.5 × 10⁵ cells in 250 µL of culture medium. Samples included unstained cells, cells incubated with trastuzumab, trastuzumab-FITC conjugate (TFC), sequentially with trastuzumab and TFC, TDCC, sequentially with TDCC and TFC. TFC, trastuzumab or TDCC was added to each sample at a final concentration of 2.5 µg/mL. Unstained cells, cells incubated with trastuzumab, TFC and TDCC were incubated for 2 h at 37 °C. For sequential staining, cells were incubated with the first reagent for 1 h at 37 °C, followed by the addition of the second reagent and further incubation for 1 h at 37 °C. Sequential incubation was performed to evaluate competitive receptor binding and potential receptor occupancy by the conjugates. After incubation, cells were centrifuged for 5 min at 300g, washed with PBS (37 °C), and resuspended in PBS for analysis. Samples were analyzed using a CytoFLEX flow cytometer (Beckman Coulter, USA). At least 10000 events were collected per sample. Flow cytometry data were analyzed using standard gating procedures, including forward and side scatter selection to exclude debris, doublet discrimination, and fluorescence thresholding based on unstained controls.

### 2.12 Microscopy

SK-BR-3 (HER2-positive) cells were detached and seeded into 24-well plates at a density of 1.5 × 10⁵ cells per well in 1 mL of culture medium. Cells were allowed to adhere overnight prior to staining. Samples included unstained cells, cells incubated with trastuzumab, FC, sequentially with trastuzumab and TFC. TFC and trastuzumab was added at a final concentration of 2.5 µg/mL. Unstained cells, cells incubated with trastuzumab and TFC were incubated for 2 h at 37 °C. For sequential staining, cells were incubated with the first reagent for 1 h at 37 °C, followed by the addition of the second reagent and further incubation for 1 h at 37 °C. After incubation, the medium containing unbound reagents was carefully removed, and cells were gently washed twice with warm PBS (37 °C). Fresh PBS (1 mL) was added prior to imaging. Cells were then analyzed using a fluorescence microscope equipped with a FITC filter set.

### 2.13 Cytotoxicity evaluation

The cytotoxicity of TDCC and copper-containing compounds was evaluated using microplate assay for cell viability. Cells at a logarithmic phase of growth were seeded (5 × 10^3^/well) into 96-well transparent flat-bottomed cell culture-treated plates (Biofil, China), left for adhesion overnight, supplemented with treatment solutions, and left for 72 h incubation at 37°C 5% CO_2_. For the treatment solution, a copper-containing compound was injected in the first step, followed by either cysteine or cell culture media in the second step. Cell culture media and cysteine alone were added to cells as the negative control. Cells were supplemented whether with resazurin or MTT for viability assessment: 1) 100 µg/ml resazurin added 6 h before the incubation ending; 2) 100 µg/ml MTT added 1.5 h before the incubation ending, supernatant aspirated, and formazan crystals dissolved in 200 µl of DMSO. Upon incubation, microplates were read at 570 nm absorbance on Tecan F50 Infinite (Tecan, Switzerland). Cell viability was calculated as the percentage of optical densities in sample wells normalized to the optical density of untreated cells (100%).

### 2.14 Clonogenic assay

A prolonged viability analysis of single MDA-MB-231 (HER2-negative) and SK-BR-3 (HER2-positive) cells treated with trastuzumab, TDCC alone and its combination with NAC was performed using a clonogenic assay. Cells were seeded at a density of 100 cells per 60-mm Petri dish and left to adhere for 48 hours. Subsequently, trastuzumab, TDCC and its combination with NAC were added, and the cells were incubated for 12 days (37°C, 5% CO₂). After the incubation, cells were fixed with ice-cold methanol for 10 minutes, stained with 1x crystal violet in 25% methanol solution for 10 minutes, and then washed with distilled water. Following drying, the results were documented using a ChemiDoc Touch (Bio-Rad, USA) in SYPRO Ruby Gel mode, and the number of formed colonies was counted.

### 2.15 Statistical analysis

Each TDCC synthesis scheme was performed at least 3 times.

Cytotoxicity assays were performed using 3 technical replicates for each concentration point. The cytotoxicity curve was approximated by the four parameter logistic curve-fit, and IC_50_ calculated at R^2^ > 0.9 using GraphPad Prism version 8.4.3 (GraphPad Software, USA).

Clonogenic assays were performed using 3 technical replicates.

Flow cytometry assay included 2-step gating approach: All events → Cells → Singlets. Each *in vitro* sample was written in 20 thousand singlets for further subpopulation markers. The abort events rate for all events was less than 5%. Raw flow cytometry data was recorded using CytExpert version 2.5 (Beckman Coulter, USA).

## 3. Results

### 3.1 DOTA-NHS ester does not hamper the redox chemistry and cytotoxicity of copper in a solution

As was shown previously [16] copper is able to participate in redox transitions in various organic contexts. However, some organic scaffolds prevent copper from interaction with NAC, so our first goal was to prove that the DOTA-NHS ester complex with copper can undergo the same transitions as previously tested compounds with similar enhancement in cytotoxicity. Another important task was to see that the DOTA-NHS ester binds copper in the solution. For this second task we used spectrophotometry and the ability of copper to precipitate phosphates in PBS (Figure 1A).

**Figure 1.**
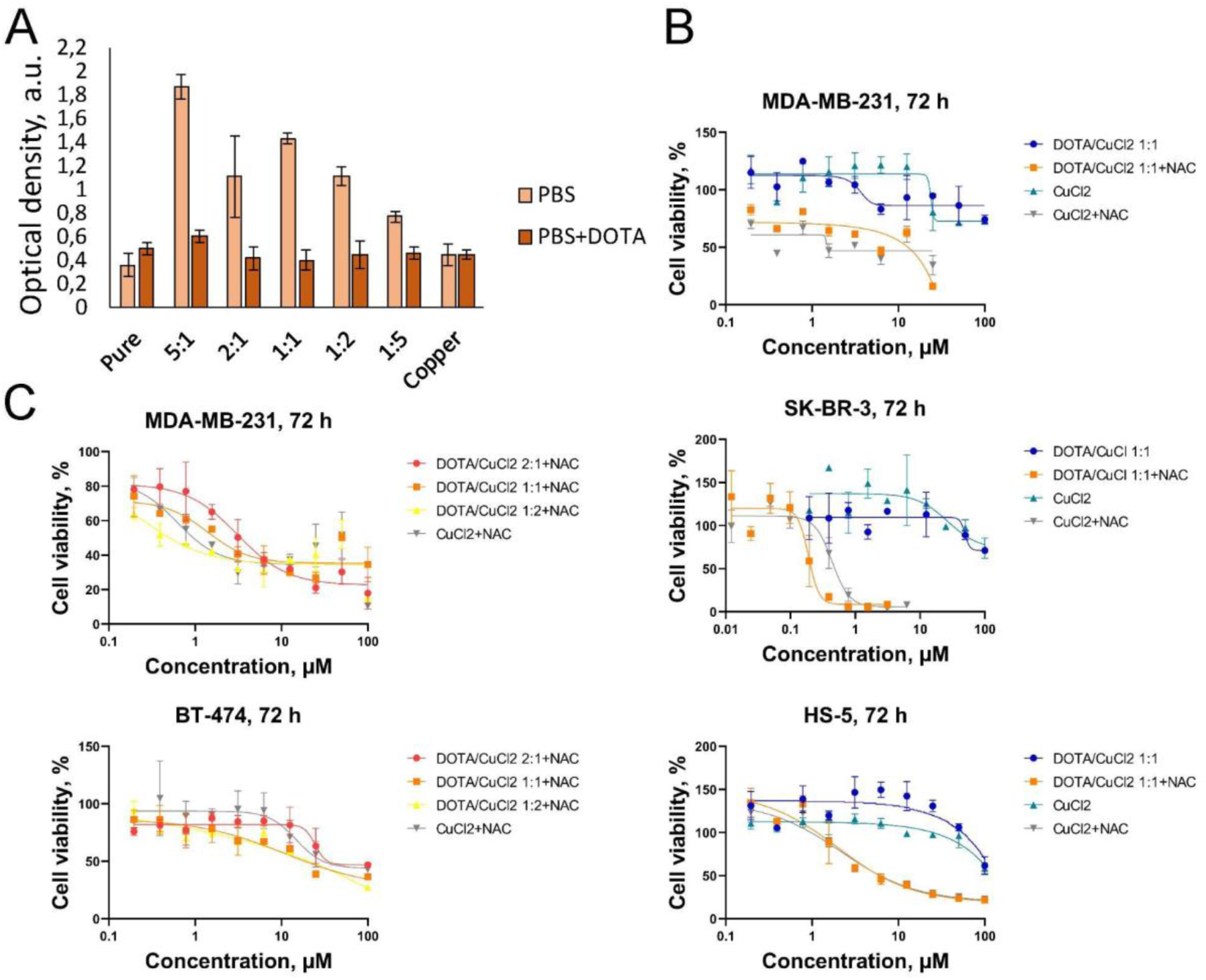
Copper-DOTA complex is able to participate in redox reactions and generate ROS upon N-acetylcysteine (NAC) or cysteine addition. A) Spectrophotometric measurement of optical density at 450 nm of solutions formed from copper-chloride and PBS or copper-acetate and PBS with DOTA. B) Cytotoxicity studies of copper-DOTA complex mixed in 1 to 1 ratio and an addition of NAC or cysteine as a reducing agent. C) MTT-assays with various proportions of DOTA.

We observed the formation of cloudy bluish precipitate upon mixing copper solution with PBS. With increased proportion of copper, the density of this precipitate decreased which can be seen on the image. However, presence of DOTA-NHS ester in the second case prevented the formation of precipitate, thus the optical density in these wells was almost the same as for the pure solutions of copper chloride or PBS (Figure 1A).

To evaluate how DOTA influences the cytotoxicity of copper in combination with NAC we performed MTT-assays. Various concentrations of CuCl_2_ were mixed with constant concentration of NAC (1 mM) and subsequent level of cell death was measured with IC_50_ values being compared for samples with or without presence of DOTA in equimolar with CuCl_2_ concentration (Figure 1B). The maximum concentration of DOTA or CuCl₂ was 100 µM for SK-BR-3, BT-474, MDA-MB-231, and HS-5 cells, followed by two–fold serial dilution. For the SK-BR-3 cell line, which appeared more sensitive to the treatment, the maximum concentration of DOTA or CuCl₂ in combination with NAC was 6.25 µM. Results for SK-BR-3, HER2-positive model, showed no difference upon DOTA addition but revealed the strongest effect of the panel upon NAC addition for both the DOTA:CuCl₂ and CuCl₂-only samples (Figure 1B, middle graph), with IC₅₀ values of 0.19 and 0.44 µM, respectively (Table 1). Similar results were observed for MDA-MB-231 and HS-5 cells: addition of DOTA-ester does not hamper the cytotoxicity of copper + NAC combination (see Table 1 for IC_50_ values).

**Table 1.**

| Cell line | $\text{IC}_{50}$ (DOTA/$\text{CuCl}_2$ (1:1) + NAC), $\mu\text{M}$ | $\text{IC}_{50}$ ($\text{CuCl}_2$ + NAC), $\mu\text{M}$ |
| --- | --- | --- |
| SK-BR-3 | 0.19±0.02 | 0.44±0.05 |
| BT-474 | 7.22±3.52 | 14.81±4.86 |
| MDA-MB-231 | 1.39±0.56 | 1.07±0.63 |
| HS-5 | 2.17±0.42 | 2.34±0.31 |

In the next experiment, different DOTA:CuCl₂ ratios (2:1, 1:1, and 1:2) were tested in order to evaluate possible cytotoxicity changes with changed ratios of the components (Figure 1C). For the HER2-positive cell line BT-474 (Figure 1C, bottom graph), a twofold excess of DOTA slightly reduced the reactivity of copper in combination with NAC. The 1:1 and 1:2 ratios showed no significant differences, whereas the sample containing only CuCl₂ with NAC did not exhibit a pronounced effect on reactivity, which is confirmed by the IC₅₀ values (11.01 µM for DOTA:CuCl₂ 1:1 versus 16.42 µM for CuCl₂). For the HER2-negative cell line MDA-MB-231 (Figure 1C, top graph), different DOTA:CuCl₂ ratios did not notably affect copper reactivity in combination with NAC.

Therefore, DOTA indeed binds copper in solution and this binding does not affect metal reactivity in combination with the reducing agent.

### 3.2 Synthesis and characterization of copper-antibody conjugates

In order to synthesize the desired copper-antibody conjugates we used conventional protocol utilized for the synthesis of radiopharmaceuticals (see Materials and methods section). DOTA is a chelator for the transition metals that can bind copper with high affinity. DOTA linkage to the antibody was performed using DOTA-NHS compound, which connects DOTA to lysine residues on the antibody. The scheme of the synthesis is shown on (Figure 2A).

**Figure 2.**
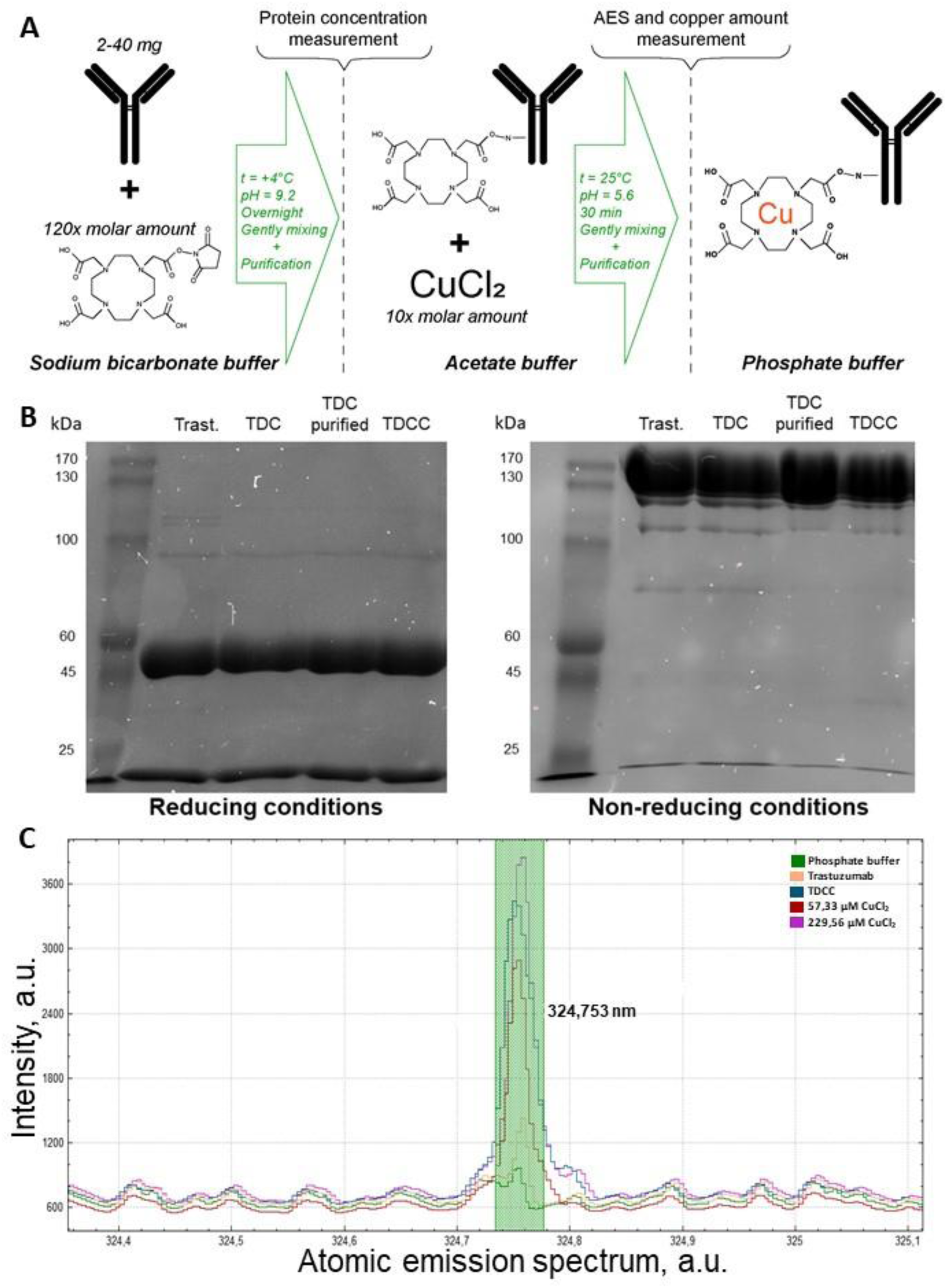
Synthesis and characterization of trastuzumab-DOTA-copper conjugate (TDCC). A) Scheme of the synthesis (for the full description see Materials and Methods section). B) SDS-PAGE of TDCC and its fractions for their integrity evaluation under reducing (with β-mercaptoethanol) and non-reducing conditions. C) Qualitative representation of copper content in TDCC compared to calibration and plain trastuzumab samples via AES.

At all stages of the synthesis the original antibody and corresponding conjugate were subjected to the evaluation of stability and level of degradation via SDS-PAGE. The SDS-PAGE analysis under reducing conditions revealed two characteristic bands for all synthesis fractions (trastuzumab, TDC before and after purification, TDCC), corresponding to molecular masses of approximately 50 kDa and 20 kDa, which represent the heavy and light chains of the antibody, respectively (Figure 2B, left image).

The SDS-PAGE analysis under non-reducing conditions showed the characteristic band for all synthesis fractions, corresponding to molecular mass of approximately 145.5 kDa, which correlates with the molecular weight of trastuzumab (Figure 2B, right image). Mentioned patterns confirm the integrity of the antibody structure and indicate that the conjugation and purification procedures did not induce degradation or aggregation of the polypeptide chains.

To evaluate the copper content in the TDCC, AES analysis was performed (Figure 2C, Figure S3). The results showed that the peak of intensity for TDCC at the 324 nm spectral line fell between calibration peaks corresponding to 229.56 μM and 57.33 μM CuCl₂. This confirms that the use of such standard samples is suitable for quantifying the TDCC concentration. Based on the measured intensity, concentration of copper was calculated, and subsequently the DAR was determined (Table 2). Moreover, the much higher peak for TDCC compared to plain trastuzumab confirms effective copper conjugation. However, due to the distribution in intensity level, caused by the tool limits, led to necessary approximations in final copper counting (Figure S3).

**Table 2.** Characteristics of obtained TDCC.

|  | **TDCC, purified with size-exclusion chromatography** | **TDCC, purified with dialysis** |
| --- | --- | --- |
| C, mg/mL<br>(by protein content) | 0.09 | 2.04 |
| C, mg/mL<br>(by copper content) | $0.63 \times 10^{-6}$ | $1.36 \times 10^{-6}$ |
| DAR<br>(copper ions per antibody) | >70.1 | 8.3±3.3 |
| Protein yield, % | 4.4 | 34.0 |

We also compared the effectiveness of purification with different methods: size-exclusion liquid chromatography and dialysis. Our results show that dialysis is more suitable for this task since it allows to obtain higher amounts of target product in smaller volumes without need for further concentration with lyophilization or other potentially detrimental for the quality of final product techniques (Table 2). Size-exclusion chromatography appeared less efficient for desalting, yielding a high DAR (>70.1), which confirms the presence of unbound copper salt in the sample solution. Moreover, changes of DAR were monitored over time (Table S1).

### 3.3 Affinity and stability of trastuzumab-DOTA-copper conjugate (TDCC)

The capability of TDCC to be stored at standard conditions of +4°C and -20°C was tested via SDS-PAGE in reducing and non-reducing conditions (with and without β-mercaptoethanol). It was performed to assess the influence of the synthesis process on the primary structure, degraded parts and dimerization on each stage of antibody modification (Figure 3). The presence of degradation pieces in the initial non-loaded trastuzumab reached 25.37% of its gel band intensity and presence of di- and polymers was 7.40% of its gel band intensity on the 10th day of its storage at +4°C (Figure 3B, -BME, +4°C, column 1), and 33.49% and 4.99% at -20°C respectively (Figure 3B, -BME, -20°C, column 1). On the 30th day of +4°C storage, non-loaded trastuzumab contained 13.31% and 14.90% of band intensity for the polymer and degraded pieces fractions respectively (Figure 3F). The distribution of the light and heavy antibody chains among all fractions and storing points was in the range from 23:71% (Figure 3D, +BME, +4°C, column 1) to 37:60% (Figure 3F, +BME, +4°C, column 1).

**Figure 3.**
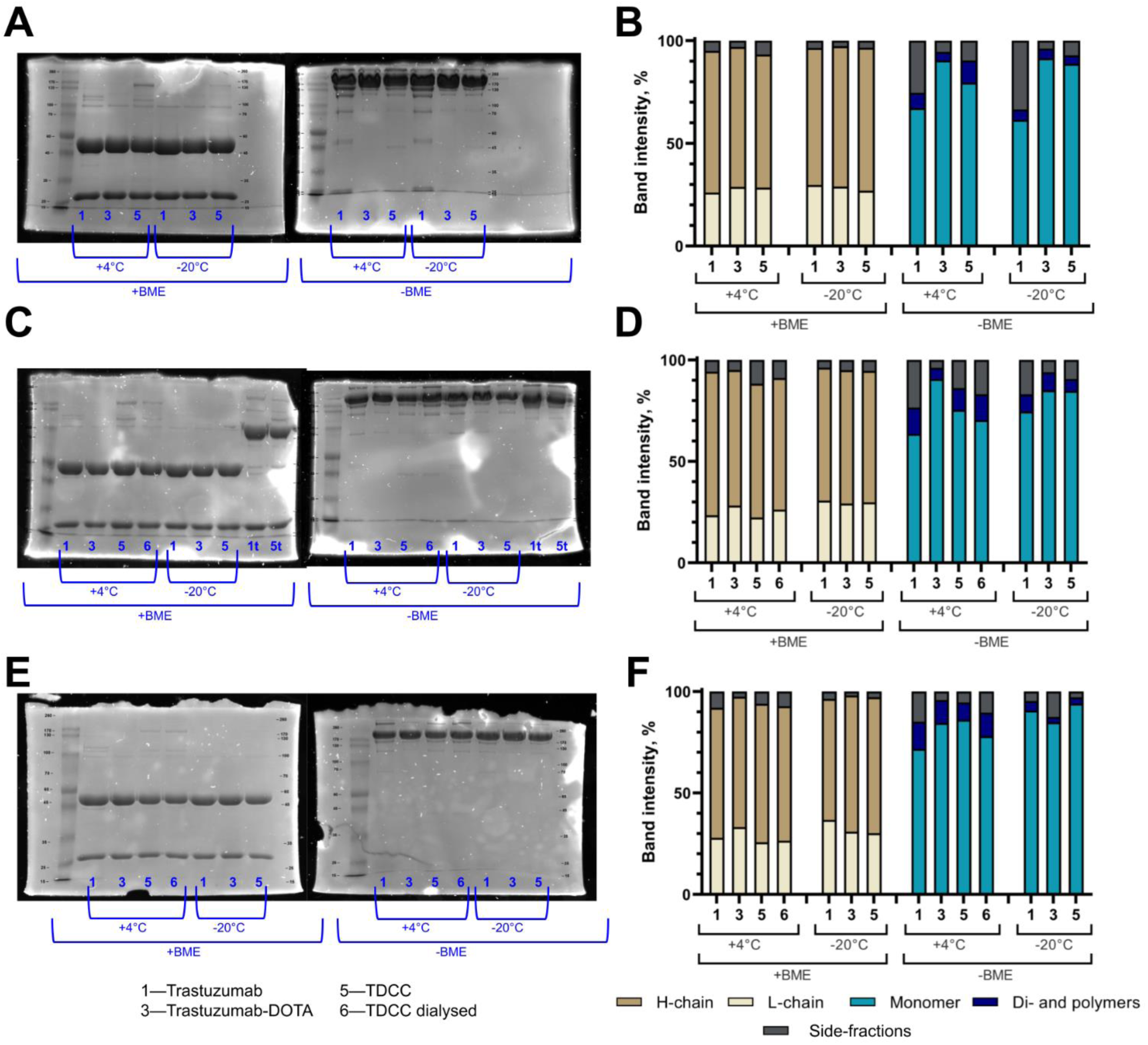
Assessment of storing conditions and synthesis procedures on the protein stability. SDS-PAGE imaging on the 10th, 20th, 30th day of storage (Figure 3A, C, E). Each gel track represents a separate sample of antibody fraction taken away during the synthesis process and stored at represented temperature and for represented time point. The - 20°C samples were frozen and thawed once for the stability assessment. Manual annotation of the gel scans (Figure 3B, D, F) including presence of light and heavy chains with partially degraded pieces (+BME groups) and presence of dimerized fraction with partially degraded pieces (-BME groups)

The affinity of synthesized TDCC was evaluated using both ELISA and competitive binding with Trastuzumab-FITC conjugate (TFC) (Figure 4). TFC was used as a convenient method for the affinity evaluation, which allows to determine the affinity toward the receptor present on the cell membrane. The results of the TFC synthesis are present in supplementary materials (Figure S4A,B). The gating strategy for flow cytometry analysis is also provided in supplementary materials (Figure S4C).

**Figure 4.**
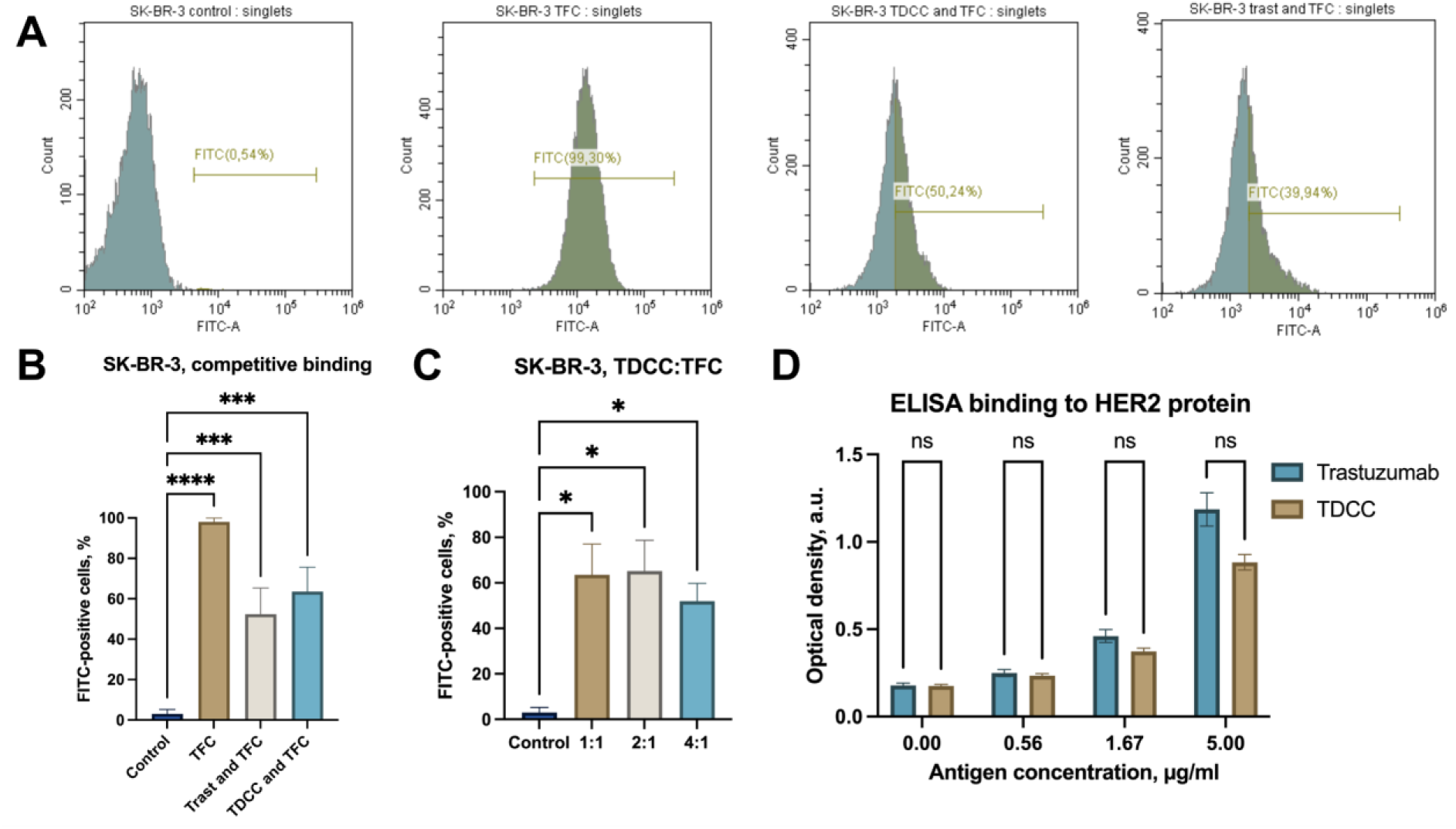
Binding properties and competitive interaction of trastusumab-FITC-conjugate (TFC) and trastuzumab-DOTA-copper conjugate (TDCC) with HER2-positive cells. A) Representative flow cytometry histograms of SK-BR-3 (HER2-positive) cells incubated under different staining conditions: unstained control, TFC, sequential incubation with TDCC followed by TFC, and sequential incubation with trastuzumab followed by TFC. B) Quantitative analysis of FITC-positive cells after incubation with TFC or after pre-incubation with trastuzumab or TDCC (n = 6). C) Quantitative analysis of FITC-positive cells after incubation with TDCC and TFC at different molar ratios (1:1, 2:1, and 4:1) (n = 3). D) ELISA analysis of TDCC and trastuzumab binding to HER2 protein to evaluate preservation of antigen recognition by the TDCC Fab fragment. Data are presented as mean ± SD from independent experiments. Statistical significance in panels B and C was determined using one-way ANOVA followed by Šídák’s multiple comparisons test. Statistical significance in panel D was determined using multiple unpaired two-tailed t-tests with Welch correction, with adjustment for multiple comparisons using the Holm-Šídák method. Differences were considered statistically significant at *p < 0.05; ***p < 0.001; ****p < 0.0001; ns, not significant.

Synthesis and purification of the TFC were assessed by gel filtration chromatography. The fraction with the highest protein concentration (832.15 ± 15.06 μg/mL) was used for subsequent binding experiments (Figure S4A,B). Spectrophotometric analysis demonstrated the degree of labeling (DOL) of 4.44 ± 0.33 FITC molecules per antibody molecule (n=6).

Competitive binding of TFC and TDCC to HER2-positive cells was evaluated using the SK-BR-3 cell line. Incubation with TFC resulted in a strong fluorescence signal, with 98.07 ± 1.90% of cells identified as FITC-positive, indicating efficient binding to HER2 receptors (Figure 4A,B). Preincubation with trastuzumab reduced subsequent binding of TFC to 52.00 ± 14.35%, confirming receptor-specific competition.

Similarly, incubation with TDCC led to decrease in FITC fluorescence intensity to 63.57 ± 13.48%, demonstrating that the conjugate retained its ability to bind HER2 receptors (Figure 4B). Increasing the proportion of TDCC resulted in FITC-positive cell percentages of 63.57 ± 13.38%, 60.00 ± 14.14%, and 51.98 ± 7.82% for TDCC:TFC ratios of 1:1, 2:1, and 4:1, respectively (Figure 4C), indicating dose-dependent competition for HER2 binding sites.

To confirm the specificity of antibody binding, the same experiments were performed using the HER2-negative MDA-MB-231 cell line. No significant fluorescence signal was detected in these cells under any staining condition (<1% FITC-positive cells), indicating minimal nonspecific binding of the conjugates (Figure S4D).

Fluorescence microscopy analysis further confirmed receptor-specific binding of the TFC to HER2-positive SK-BR-3 cells. Strong membrane-associated fluorescence was observed in cells incubated with FC, whereas minimal signal was detected in untreated control cells and in cells incubated with trastuzumab. Pre-incubation with trastuzumab followed by TFC resulted in a reduced fluorescence signal, consistent with competitive receptor binding (Fig S4 E).

ELISA analysis was performed in two independent experiments and demonstrated comparable binding of TDCC and intact trastuzumab to HER2 protein across the tested antigen concentration range (Figure 4D). Both antibodies exhibited a concentration-dependent increase in optical density values, indicating preserved antigen recognition by the TDCC Fab fragment. At the highest HER2 concentration (5.0 µg/mL), optical density values reached 1.19 for trastuzumab and 0.88 for TDCC, consistent with slightly reduced but maintained binding activity of the conjugate. These results indicate that conjugation did not substantially impair HER2-binding capability of the TDCC.

### 3.4 Cellular effects of prolonged treatment with TDCC

TDCC is able to reduce viability of both HER2-positive and triple-negative resistant breast cancer cells.

In the clonogenic assay the colony formation inhibition potential of synthesized TDCC was observed (Figure 5). Trastuzumab and TDCC were used at a concentration of 5 μg/mL (by protein content), while NAC was used at 1 mM.

**Figure 5.**
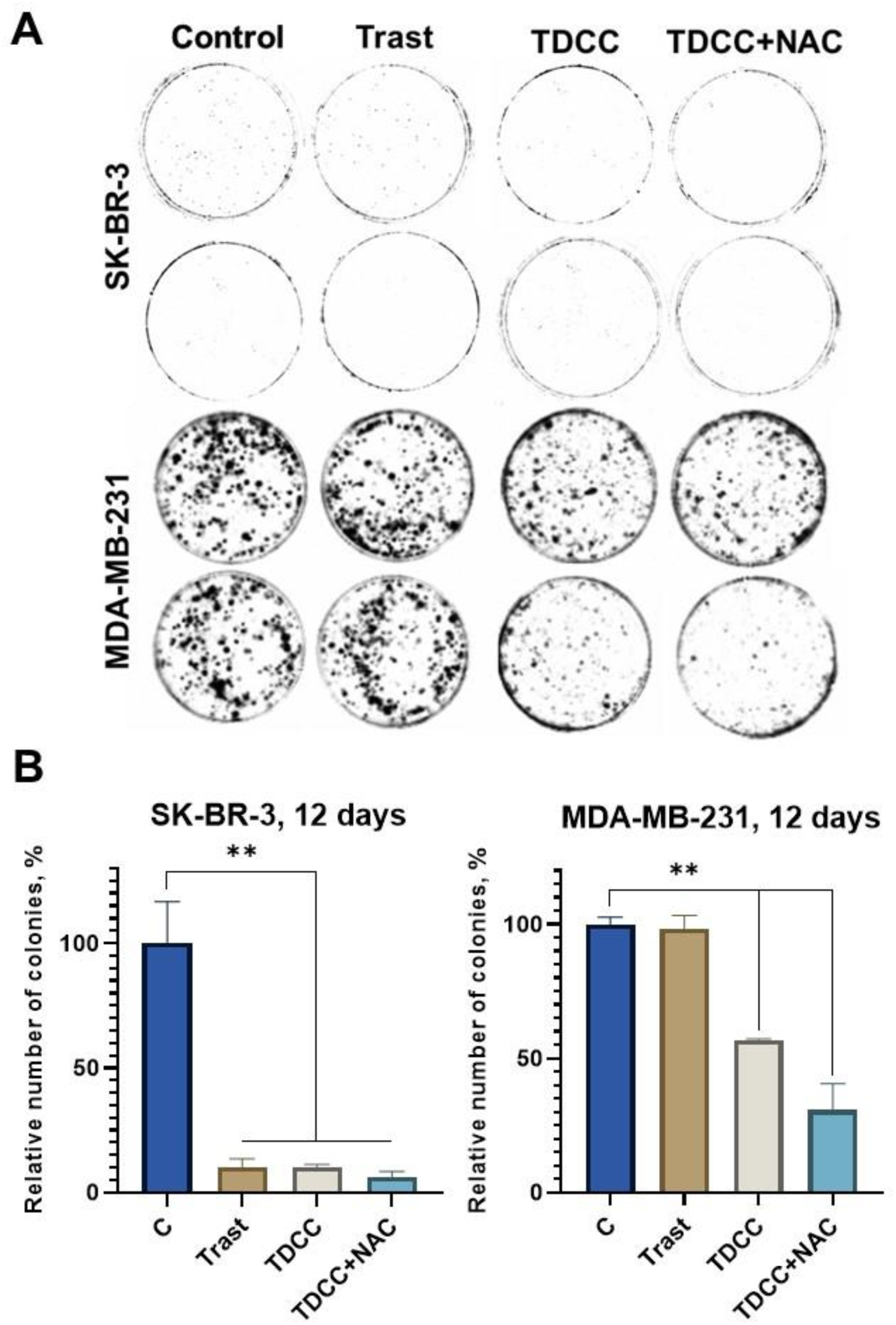
Colonies formation of MDA-MB-231 and SK-BR-3 after treatment with trastuzumab (Trast), trastusumab-DOTA-copper conjugate (TDCC), and TDCC in combination with N-acetylcysteine (NAC). A) Representative images of colonies formed in culture dishes under different treatment conditions. B) Quantification of colony numbers in histograms (**p < 0.01).

In HER2-positive SK-BR-3 cells, trastuzumab, TDCC, and TDCC with NAC each caused a considerable reduction in the relative number of colonies compared to the control, with the most pronounced effect observed for the TDCC and NAC combination (Figure 5B, left histogram).

In contrast, HER2-negative MDA-MB-231 cells showed no significant change in colony formation upon trastuzumab treatment relative to the control. However, both TDCC alone and TDCC with NAC inhibited colony formation on this model, with the combination again demonstrating the most pronounced effect (Figure 5B, right histogram). The reduction of cell viability of MDA-MB-231 (HER2-negative) cells can be attributed to the long-term effect of prolonged incubation with the combination of TDCC and NAC, which results in accumulation of cell damage. Thus, both TDCC and TDCC in combination with NAC inhibit colonies’ formation on HER2-positive and HER2-negative cellular models.

### 3.5 TDCC causes ROS formation *in vitro*

Cytotoxicity of TDCC and its combination with NAC was evaluated via the MTT assay in comparison to CuCl2 treatment in SK-BR-3 and MDA-MB-231 cells (Figure 6A). The maximum concentration of CuCl₂ was 1.05 µM, and for TDCC - 1.1 µM by protein and copper content, respectively. The concentration of NAC was fixed at 1 mM. In HER2–positive SK–BR–3 cells, both the combination of CuCl₂ with NAC and TDCC with NAC decreased cell viability, with IC_50_ values of 0.5215 ±0.2152 μM for TDCC + NAC and 0.3769 ±0.1457 μM for CuCl2+NAC. On the other hand, no such pronounced effect was observed in MDA-MB-231 cells.

**Figure 6.**
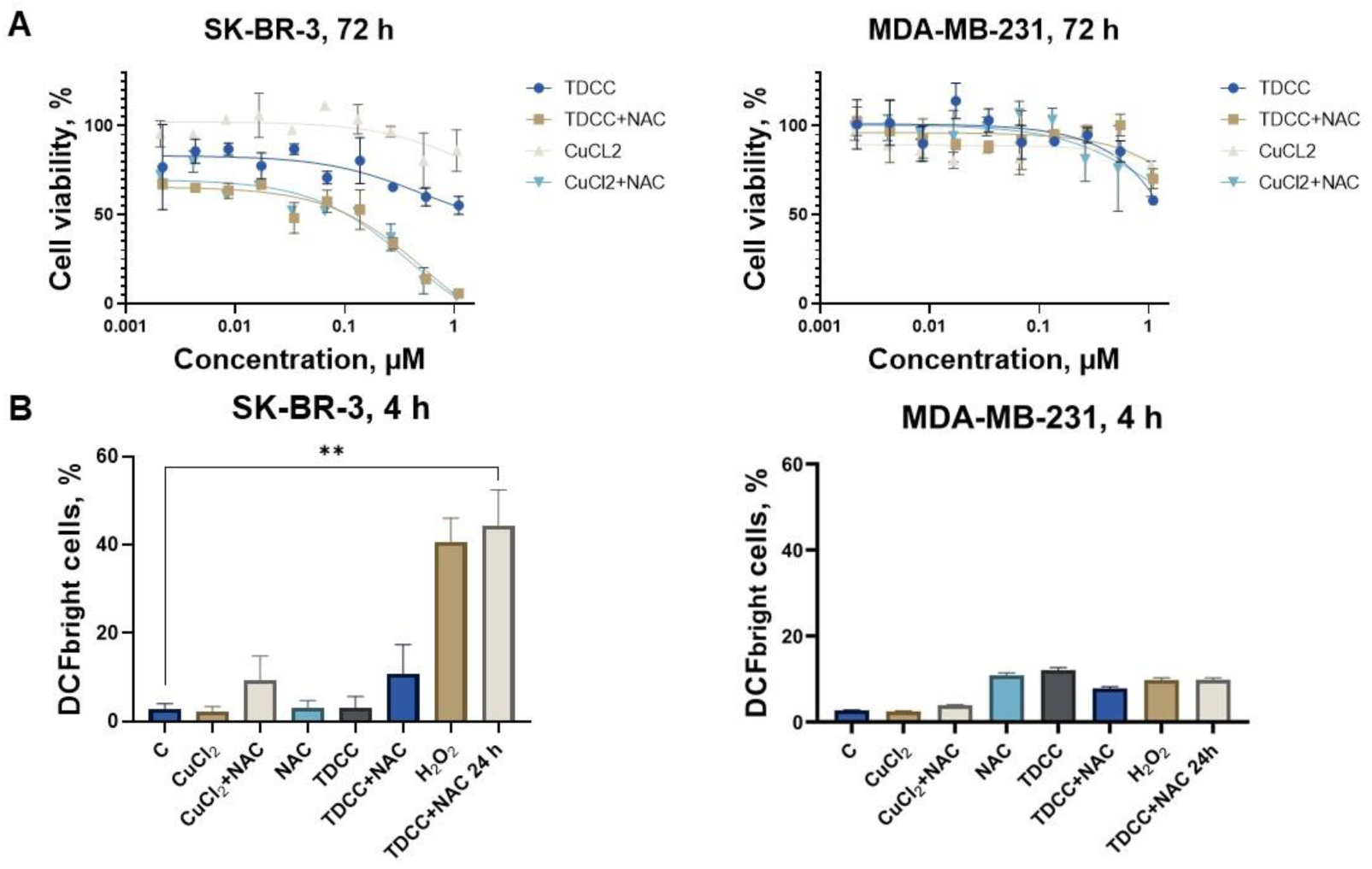
ROS accumulation and cytotoxicity under the treatment of CuCl₂ or trastuzumab-DOTA-copper conjugate (TDCC) (alone or in combination with N-acetylcysteine (NAC)). A) Cytotoxicity in SK-BR-3 and MDA-MB-231 cells B) ROS accumulation in SK-BR-3 and MDA-MB-231 cells (**p < 0.01).

ROS accumulation in cells upon TDCC treatment, compared to CuCl₂ treatment, was assessed using the H_2_DCFDA dye by flow cytofluorimetry, with the fluorescent signal detected in the FITC channel (Figure 6B, Figure S5). CuCl₂ and TDCC were used at a concentration of 5 μM (by copper content for TDCC), while NAC was used at 1 mM. Treatment lasted for 4 hours. Additionally, the TDCC+NAC sample was also incubated for 24 hours to assess maximum ROS accumulation. H₂O₂ was used at 1 mM for 1-2 hours as a positive control.

In HER2-positive SK-BR-3 cells, both the combination of CuCl₂ with NAC and TDCC with NAC led to a slightly increased intensity of DCFDA fluorescence compared to the negative control, confirming ROS accumulation. The maximum ROS accumulation was observed in the TDCC+NAC sample at 24 hours, which correlated with ROS level in the positive control sample (H₂O₂).

On the other hand, in HER2-negative MDA-MB-231 cells, no such pronounced increase (p > 0.05 in ROS levels was observed for either the TDCC+NAC (24 hours) or the positive control (H₂O₂). Only a slight increase in ROS levels was detected in the samples treated with NAC alone, TDCC alone, TDCC+NAC (4 hours), TDCC+NAC (24 hours), and H₂O₂. This can be attributed to the enhanced activation of antioxidant systems in these cells [30,31].

Thereby, the combination of TDCC with NAC leads to ROS accumulation, with a pronounced effect on the HER2-positive SK-BR-3 cell line.

## 4. Discussion

Nowadays ADCs are advancing in the field of cancer therapy with around 15 compounds being approved by FDA and 20 compounds globally [32] for various malignancies. However, major drawbacks of ADCs still hamper their development: preliminary payload release due to linker cleavage (which triggers side effects), extremely high cytotoxicity of payloads (which requires special equipment for the production of ADC increasing costs), low versatility of cell death mechanisms caused by the payload (which can result in additional drug resistance and lower drug effectiveness). These factors cause constant search not only for new payloads but also linkers and structures that can enhance ADCs effectiveness. We believe that one of these advancements could be the use of chemodynamic approaches, described previously [33,34] and also in our previous work [16].

Thus, this work shows the synthesis and *in vitro* efficacy assessment of novel TDCC, a type of ADC with active transition metal payload able to generate ROS in reducing conditions. The concept is based on the combination of copper and NAC, which was shown to be highly cytotoxic against various cell lines. However, this effect is non-specific, meaning that it can damage healthy tissues and organs. To mitigate that obstacle, we developed the design of antibody conjugates where copper can serve as a payload and exert its action only in the presence of elevated levels of NAC. ROS generated by this interaction can damage the cell membrane of resistant cells as well as surrounding cells of the tumor microenvironment. Because the process does not rely on internalisation or drug efflux pumps, it may bypass these resistance mechanisms, although upregulation of antioxidant defences could potentially limit its efficacy [35]. This approach combines several modalities: non-internalizing ADC, chemodynamic influence, and combination therapy.

We chose trastuzumab, an antibody against HER2 receptor which is often overexpressed on breast cancer cells, for modification. A conventional DOTA-NHS ester, which is used for the production of radiopharmaceuticals, was used for copper conjugation. While classical ADCs often utilize cleavable linkers, our design implies that a stable complex of DOTA and copper persist until the conjugate reaches the tumor. DOTA chelator provides high kinetic stability: copper is not released before reduction, and the cytotoxic activity is triggered only when Cu²⁺ is reduced to Cu⁺. The successful synthesis of trastuzumab-DOTA-copper conjugate (TDCC) was confirmed by multiple methods, which indicate that conjugation reaction does not hamper the structure of antibodies, final DAR reaches approximately 8.3, affinity to HER2 remains intact. Biological activity of TDCC allows the use of it for the treatment of HER2-positive malignancies, but prolonged exposure can also be harmful for HER2-negative cells, which brings additional promises about overcoming drug resistance and concerns about safety issues.

Future experiments will be directed toward additional characterization of cellular and molecular mechanisms of action of TDCC, detailed description of chemical redox reactions in the system, and pre-clinical studies on animals. Furthermore, many issues also require additional attention, e.g. DAR stabilization and increase, possibility of using other linkers and chelators, stability and safety under prolonged times of treatment and in physiological conditions. These aspects are important considerations that may arise with the progress of drug development; however, the purpose of current work was to show that the antibody-copper conjugates can be synthesized and that they have the desired activity against targeted cancer cells. This finding gives a way for many possibilities of modification and optimization for better outcomes.

## Acknowledgements

This work is an output of a research project (HSE-BR-2025-76) implemented as part of the Basic Research Program at HSE University.

## Author contributions

D.O., K.Ch., M.M, and S.G. performed experiments. D.O., K.Ch., S.G., and S.T. statistical analysis and visualization. D.O., K.Ch., O.K., and S.T. writing the original draft. S.T., V. Sh., and O.K. conceptualization and study design. S.T. and V.Sh. supervision.

## Data availability

Data will be made available on request.

## Abbreviations

Ab: antibody
ADC: antibody–drug conjugate
DAR: drug-to-antibody ratio
FITC: fluorescein isothiocyanate
HER2: human epidermal growth factor receptor 2
NAC: N-acetylcysteine
TDC: trastuzumab-DOTA conjugate
TDCC: trastuzumab-DOTA-copper conjugate
TFC: trastuzumab-FITC conjugate
SEC: size exclusion chromatography.

