## Supplementary Materials for "Synthesis and evaluation of novel copper-antibody conjugates for the chemodynamic therapy of HER2-positive breast cancer"

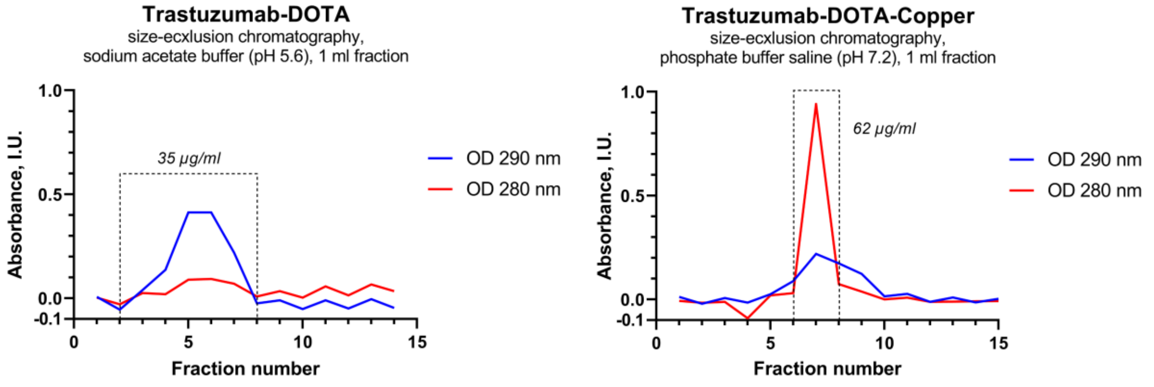


**Figure S1. The results of gel-filtration chromatography for the first stage of TDCC synthesis**

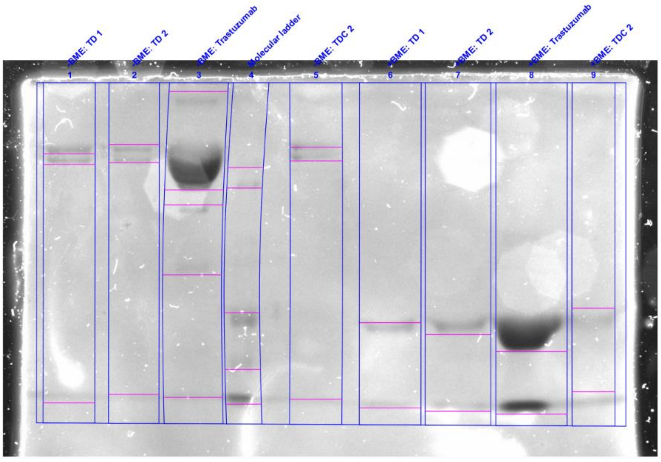


**Lane 1 - -BME: TD 1**

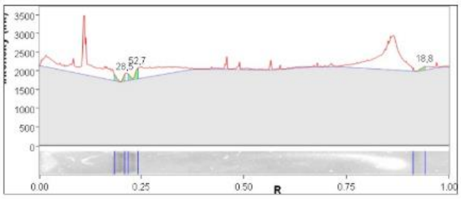


| Band No. | Band Label | Mol. Wt. (KDa) | Relative Front | Adj. Volume (Int) | Volume (Int) | Abs. Quant. (mg) | Rel. Quant. | Band % | Lane % |
|---|---|---|---|---|---|---|---|---|---|
| 1 | | N/A | 0,209 | 323 456 | 7 598 328 | N/A | N/A | 28,5 | 1,3 |
| 2 | | N/A | 0,240 | 597 968 | 8 076 064 | N/A | N/A | 52,7 | 2,5 |
| 3 | | N/A | 0,941 | 213 560 | 9 985 184 | N/A | N/A | 18,8 | 0,9 |

| | |
|---|---|
| Lane Background | Lane background subtracted with disk size: 10 |
| Lane Width | 7.28 mm |

**Figure S2. Gel annotation in ImageLab and the values of intensity for chosen bends in the first line**

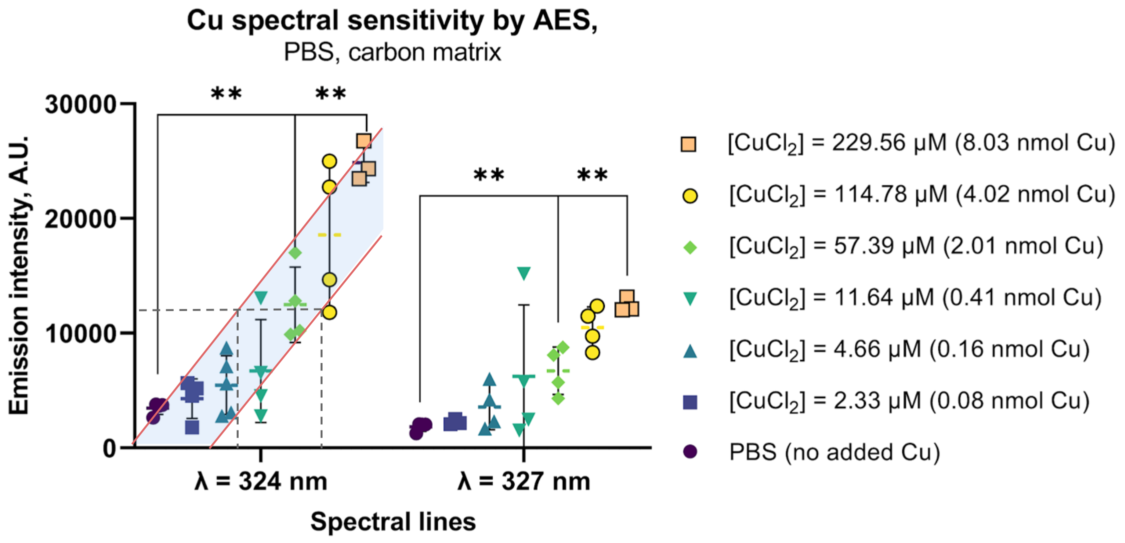


**Figure S3. AES (Atomic Emission Spectroscopy).** Distribution of absorption intensity of 327 nm and 324 nm for the samples with known copper concentration. According to the distribution in control samples the calibration curves were built allowing to determine the range of DAR values.

*Table S1. Time-dependent DAR alterations*

| | **1-3 days** | **10 days** | **20 days** |
|---|---|---|---|
| DAR min | 5.3 | 0.6 | 0.5 |
| DAR max | 11.3 | 3.4 | 3.9 |

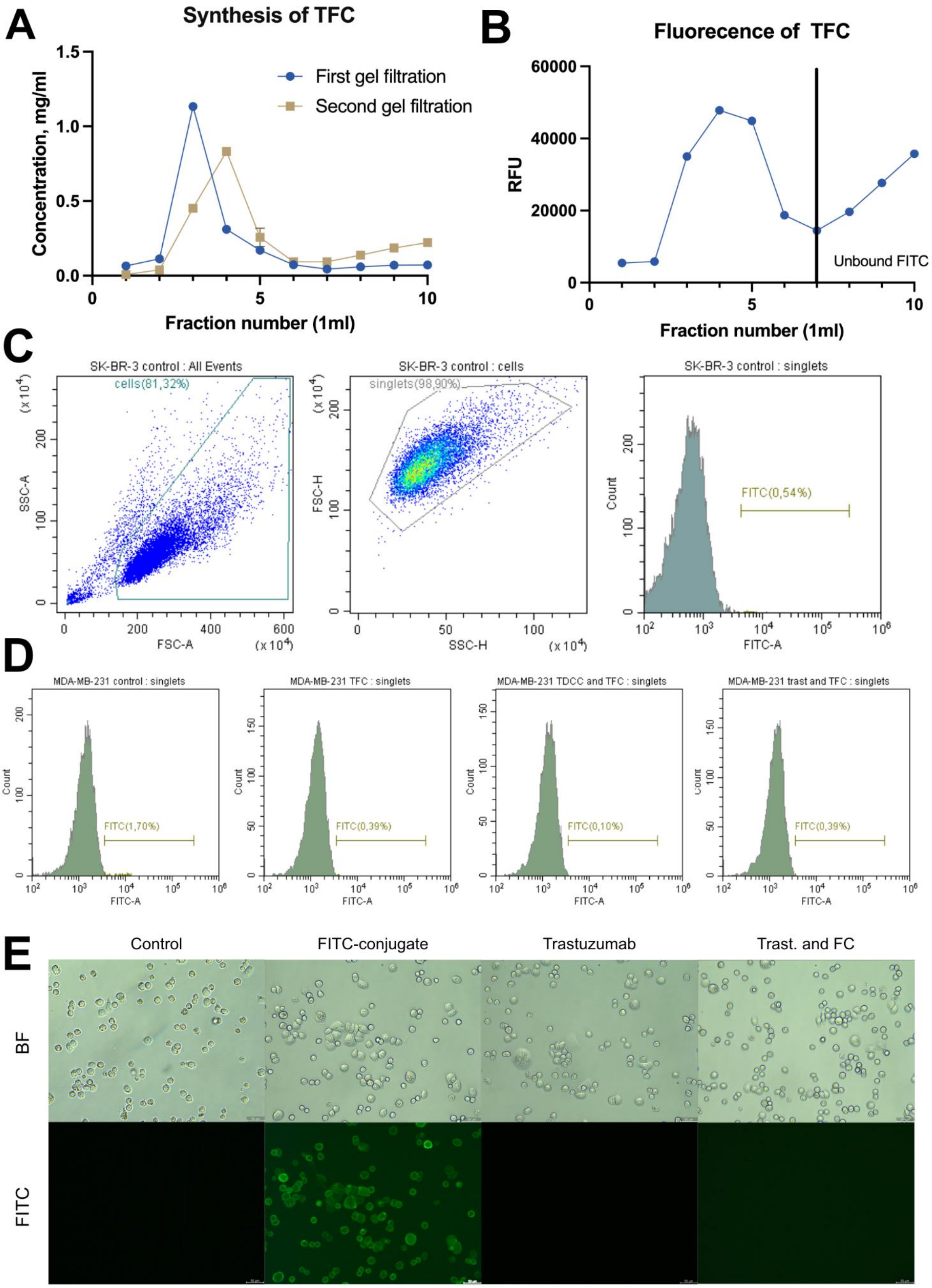


**Figure S4. Purification, characterization, and receptor-specific binding analysis of the trastuzumab-FITC conjugate (TFC).** A) Protein concentration profile of fractions collected during the first and second gel filtration steps used for buffer exchange and removal of unbound FITC. B) Fluorescence intensity (relative fluorescence units, RFU) of collected fractions measured at 495 nm. Fractions collected after the vertical line correspond to unbound FITC. C) Flow cytometry gating strategy used for the analysis of FITC-positive cells. Sequential gating included selection of all events, cell population based on forward and side scatter parameters, singlet discrimination, and identification of FITC-positive cells. D) Flow cytometry analysis of the HER2-negative MDA-MB-231 cell line following incubation with TFC, TDCC, or sequential combinations of these reagents under the same experimental conditions as described for the SK-BR-3 cell line. E) Fluorescence microscopy analysis of the HER2-positive SK-BR-3 cell line incubated under different staining conditions. Representative bright-field (BF) and fluorescence (FITC) images of cells incubated with control (no reagents), TFC, trastuzumab, and sequential incubation with trastuzumab followed by TFC, demonstrating receptor-specific competitive binding.

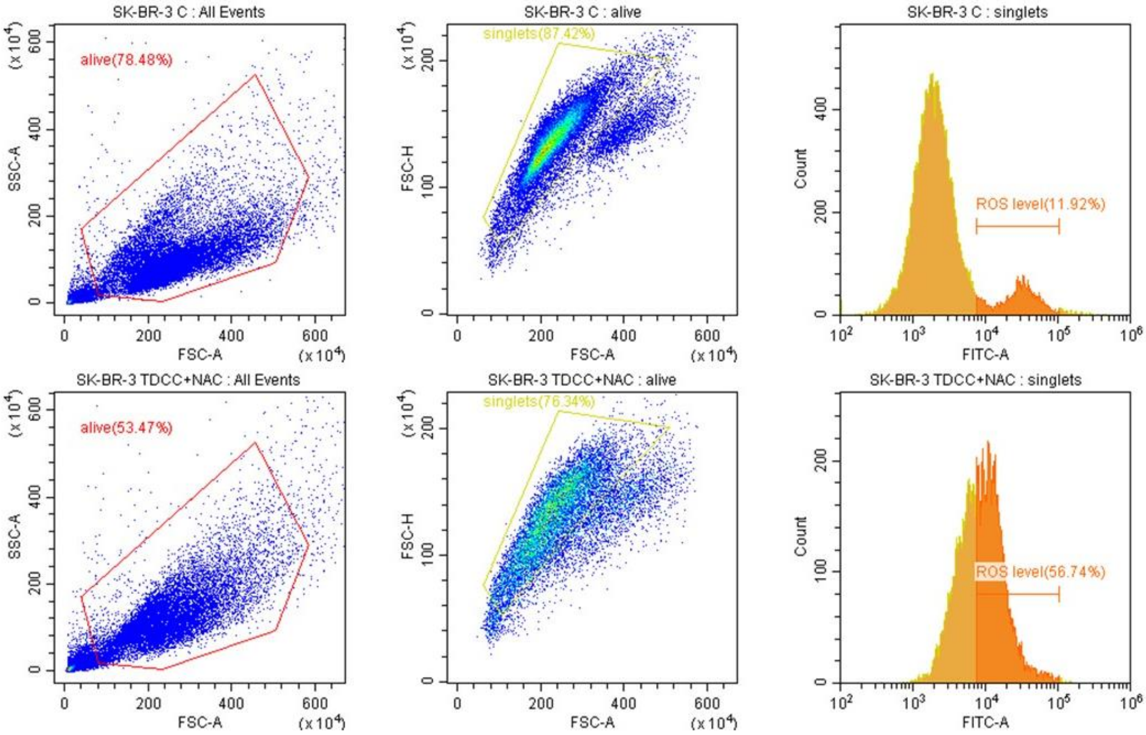


**Figure S5. Representative gating of cells stained with $H_2$DCFDA for flow cytofluorimetric ROS analysis.**
